# The SARS-CoV-2 Spike protein has a broad tropism for mammalian ACE2 proteins

**DOI:** 10.1101/2020.06.17.156471

**Authors:** Carina Conceicao, Nazia Thakur, Stacey Human, James T. Kelly, Leanne Logan, Dagmara Bialy, Sushant Bhat, Phoebe Stevenson-Leggett, Adrian K. Zagrajek, Philippa Hollinghurst, Michal Varga, Christina Tsirigoti, John A. Hammond, Helena J. Maier, Erica Bickerton, Holly Shelton, Isabelle Dietrich, Stephen C. Graham, Dalan Bailey

## Abstract

SARS-CoV-2 emerged in late 2019, leading to the COVID-19 pandemic that continues to cause significant global mortality in human populations. Given its sequence similarity to SARS-CoV, as well as related coronaviruses circulating in bats, SARS-CoV-2 is thought to have originated in Chiroptera species in China. However, whether the virus spread directly to humans or through an intermediate host is currently unclear, as is the potential for this virus to infect companion animals, livestock and wildlife that could act as viral reservoirs. Using a combination of surrogate entry assays and live virus we demonstrate that, in addition to human ACE2, the Spike glycoprotein of SARS-CoV-2 has a broad host tropism for mammalian ACE2 receptors, despite divergence in the amino acids at the Spike receptor binding site on these proteins. Of the twenty-two different hosts we investigated, ACE2 proteins from dog, cat and rabbit were the most permissive to SARS-CoV-2, while bat and bird ACE2 proteins were the least efficiently used receptors. The absence of a significant tropism for any of the three genetically distinct bat ACE2 proteins we examined indicates that SARS-CoV-2 receptor usage likely shifted during zoonotic transmission from bats into people, possibly in an intermediate reservoir. Interestingly, while SARS-CoV-2 pseudoparticle entry was inefficient in cells bearing the ACE2 receptor from bats or birds the live virus was still able to enter these cells, albeit with markedly lower efficiency. The apparently broad tropism of SARS-CoV-2 at the point of viral entry confirms the potential risk of infection to a wide range of companion animals, livestock and wildlife.

## Introduction

The β-coronavirus SARS-CoV-2 emerged in late 2019, causing a large epidemic of respiratory disease in the Hubei province of China, centred in the city of Wuhan [1]. Subsequent international spread has led to an ongoing global pandemic, currently responsible for 8 million infections and over 435,000 deaths (as of 11^th^ June 2020, John Hopkins University statistics; https://coronavirus.jhu.edu/map.html). As for SARS-CoV, which emerged in China in late 2002, and MERS-CoV, which emerged in Saudi Arabia in 2012, the original animal reservoir of zoonotic coronaviruses is thought to be bats [2]. Spill-over into humans is suspected or proven to be facilitated through an intermediate host, e.g. civets for SARS-CoV [2] or camels for MERS-CoV [3]. For SARS-CoV-2, a bat origin is supported by the 2013 identification of a related coronavirus RaTG13 from *Rhinolophus affinis* (intermediate horseshoe bat), which is 96% identical at the genome level to SARS-CoV-2 [1]. Identifying the animal reservoir of SARS-CoV-2, and any intermediate hosts via which the virus ultimately spread to humans, may help to understand how, where and when this virus spilled over into people. This information could be vital in identifying future risk and preventing subsequent outbreaks of both related and unrelated viruses. Concurrent to this, there is also a need to understand the broader host tropism of SARS-CoV-2 beyond its established human host, in order to forewarn or prevent so-called reverse zoonoses, e.g. the infection of livestock or companion animals. The latter could have serious implications for disease control in humans and consequently impact on animal health and food security as we seek to control the COVID-19 pandemic.

The process of viral transmission is complex and governed by a range of factors that in combination determine the likelihood of successful infection and onward spread. The first barrier that viruses must overcome to infect a new host, whether that be typical (of the same species as the currently infected host) or atypical (a new species) is entry into the host cell. Entry is governed by two opposing variables; the first being efficient virus binding to the host cell and the second being host-mediated inhibition of this process, e.g. through virus-specific neutralising antibodies. In the case of SARS-CoV-2, it is likely that in late 2019 the entire global population was immunologically naïve to this virus, although there is debate as to whether pre-existing immunity to the endemic human-tropic coronaviruses, e.g. OC43 and HKU1, provides any cross-protective antibodies to help mitigate disease symptoms [4]. To compound this, the rapid global spread of SARS-CoV-2, combined with emerging molecular data [5, 6], have clearly demonstrated that SARS-CoV-2 is efficient at binding to and entering human cells. However, how widely this host-range or receptor tropism extends and the molecular factors defining atypical transmission to non-human hosts remain the subject of intense investigation.

Coronavirus entry into host cells is initiated by direct protein-protein interactions between the virally encoded homo-trimeric Spike protein, a class I transmembrane fusion protein found embedded in the virion envelope, and proteinaceous receptors or sugars on the surface of host cells [7]. The high molecular similarity of β-coronaviruses, specifically SARS-CoV, allowed the rapid identification of angiotensin-converting enzyme 2 (ACE2) as the proteinaceous receptor for SARS-CoV-2 [8, 9] and structural studies characterising Spike bound to ACE2 have quickly followed [5, 6, 10, 11]. These studies have identified a high affinity interaction between the receptor binding domain (RBD) of Spike and the N-terminal peptidase domain of ACE2, which for SARS-CoV has been shown to determine the potential for cross-species infection and ultimately, pathogenesis [12].

The availability of ACE2 gene sequences from a range of animal species enables the study of receptor tropism of SARS-CoV-2 Spike. This can be used to predict whether receptor usage is likely to be a driving factor in defining the host range of this virus, either through computational predictions based on ACE2 sequence conservation [13] or, more directly, with functional experimental investigation [1]. In this paper, we examined whether ACE2 from 22 different species of livestock, companion animals and/or wildlife could support the entry of SARS-CoV-2, alongside human ACE2. Using two distinct assays we identified that SARS-CoV-2 has a broad receptor tropism for mammalian ACE2 proteins including those from hamster, pig and rabbit. Efficient infection via these ACE2 receptors was subsequently confirmed using live SARS-CoV-2 virus. Interestingly, receptors that were unable to support particle entry in pseudotype assays, e.g. chicken ACE2, were still able to support live virus entry at a high multiplicity of infection. This research has identified vertebrate species where cell entry is most efficient, allowing prioritisation of *in vivo* challenge studies to assess disease susceptibility. Combining this with increased surveillance and improved molecular diagnostics could help to prevent future reverse zoonoses.

## Results

### The SARS-CoV-2 binding site on ACE2 is highly variable

Recent structural and functional data have shown that SARS-CoV, SARS-CoV-2 and other β-coronavirus (lineage B clade 1) Spike proteins bind the same domain in ACE2 to initiate viral entry [5, 6, 8–10]. We thus hypothesised that SARS-CoV-2 could use the ACE2 receptor to infect a range of non-human, non-bat hosts. To this end we synthesised expression constructs for human ACE2 as well as orthologues from 22 other vertebrate species, including nine companion animals (dogs, cats, rabbits, guinea pigs, hamsters, horses, rats, ferrets, chinchilla), seven livestock species (chickens, cattle, sheep, goats, pigs, turkeys, buffalo), four bat species (horseshoe bat, fruit bat, little brown bat and flying fox bat), and two species confirmed or suspected to be associated with previous coronavirus outbreaks (civet and pangolin). There is 62 to 99% sequence identity between these proteins at the amino acid level (76-99% when excluding the two bird sequences) and their phylogenetic relationships are largely consistent with vertebrate phylogeny, although the guinea pig sequence was more divergent than predicted (**Fig.1A**). Examining the conservation of amino acids at the SARS-CoV-2 binding site on the surface of the ACE2 protein revealed a high degree of variation across mammalian taxa (**Fig.1B,C**), suggesting that SARS-CoV-2 receptor binding may vary between potential hosts. This variation was also evident when aligning the 23 ACE2 sequences included in our study, which identified a number of highly variable residues within the overlapping SARS-CoV and SARS-CoV-2 binding sites, including Q24, D30, K31, H34, L79 and G354 (**Fig.1D**). Our first step was to ensure efficient and equivalent surface expression of these ACE2 proteins on target cells. To this end their N-terminal signal peptides were replaced with a single sequence from the commercially available pDISPLAY construct (**Fig.1E**). In addition, the ectodomain was fused with a HA-epitope tag to allow the specific detection of surface expressed protein. Western blot of whole cell lysates together with flow cytometric analysis of cell surface expression confirmed that in the majority of cases the 23 ACE2 proteins were expressed to similar levels, thereby allowing side-by-side comparison of their usage by SARS-CoV-2 (**Fig.1F,G; Sup.Fig.1**). The marked exceptions were flying fox bat and guinea pig ACE2 (**Fig.1F,G**) where protein expression and cell-surface presentation were barely detectable. The cause of this poor expression is unknown, potentially arising due to errors in the ACE2 sequences available for these species. Since the available sequence accuracy for these two genes would need to be explored further these two ACE2 proteins were excluded from our subsequent experiments.

**Figure 1:**
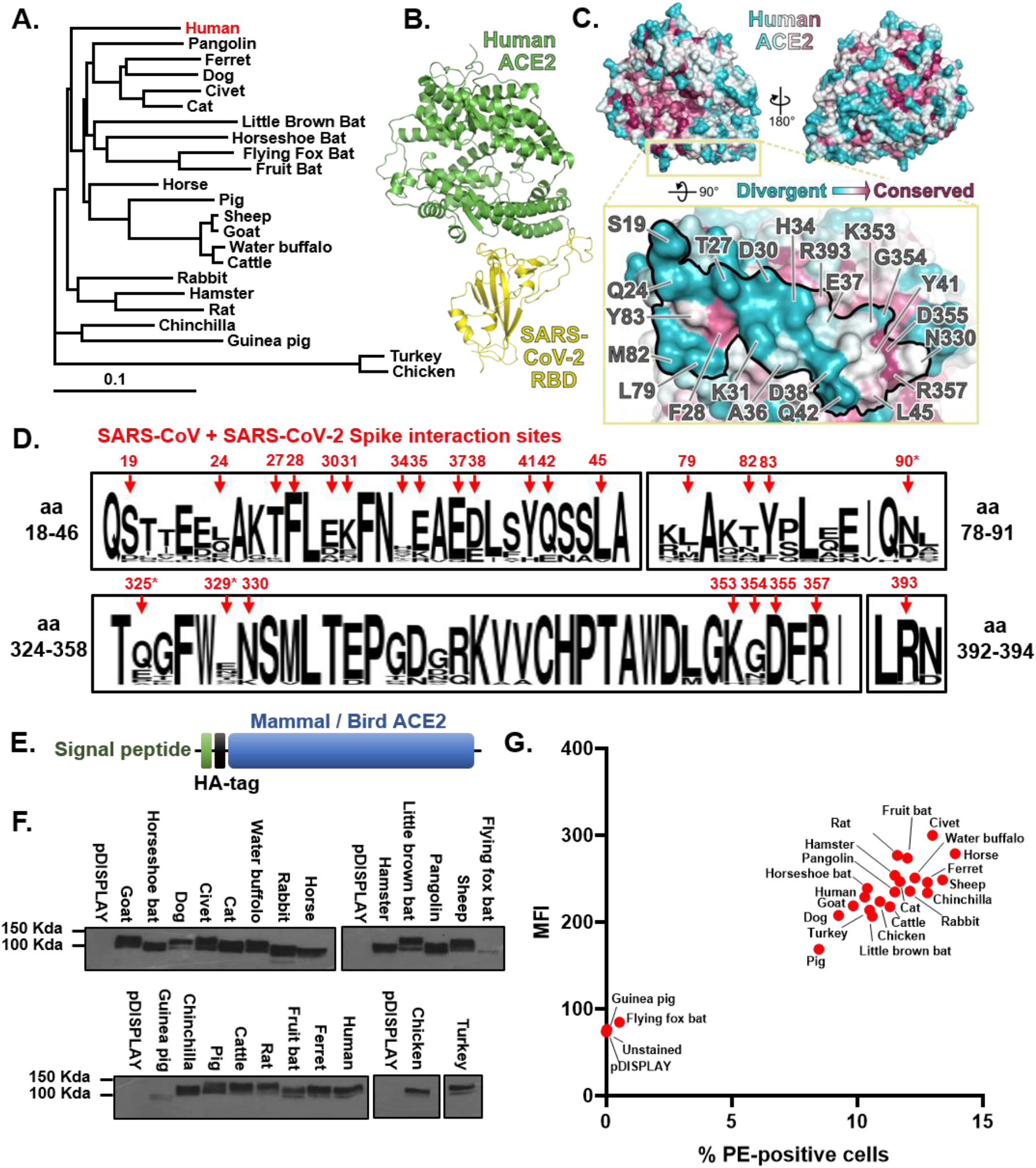
The SARS-CoV-2 binding site on ACE2 is highly variable. **(A)** A phylogenetic tree of ACE2 proteins assembled using the Neighbor-Joining method [45] conducted in MEGA7 [46] with ambiguous positions removed. The tree is drawn to scale and support was provided with 500 bootstraps. **(B)** Structure of human ACE2 ectodomain (green) in complex with the receptor binding domain (RBD) of SARS-CoV-2 [10]. **(C)** Conservation of mammalian ACE2 amino acid residues mapped onto the surface of the ACE2 ectodomain [10], coloured from blue (divergent) to purple (conserved) and two orientations. Inset depicts the SARS-CoV-2 binding region of ACE2 (outlined), with residues that contact the SARS-CoV-2 RBD highlighted [6]. **(D)** WebLogo [47] plots summarising the amino acid divergence within the mammalian and bird ACE2 sequences characterised in this study. The single letter amino acid (aa) code is used with the vertical height of the amino acid representing its prevalence at each position in the polypeptide (aa 18-46, 78-91, 324-358 and 392-394 are indicated). The aa sites bound by SARS-CoV and SARS-CoV-2 Spike [11] are indicated by red arrows. **(E)** ACE2 sequences were cloned into the pDISPLAY expression construct in frame with an N-terminal signal peptide (the murine Ig κ-chain leader sequence) and HA-tag. **(F)** Expression of individual mammal or bird ACE2 proteins was confirmed at a whole cell level by western blot. **(G)** Flow cytometry was performed to examine surface expression of each ACE2 protein on non-permeabilised cells. For gated cells the percentage positivity and mean fluorescence intensity (MFI) are plotted.

### Receptor screening using surrogate entry assays identifies SARS-CoV-2 Spike as a pan-tropic viral attachment protein

To examine the capacity of SARS-CoV-2 to enter cells bearing different ACE2 proteins we used two related approaches. The first, based on the widely employed pseudotyping of lentiviral particles with SARS-CoV-2 Spike [9], mimics particle entry. The second approach, based on a quantitative cell-cell fusion assay we routinely employ for the morbilliviruses [14], assesses the capacity of Spike to induce cell-cell fusion following receptor engagement. In both assays we used a codon-optimised SARS-CoV-2 Spike expression construct as the fusogen, demonstrating robust and sensitive detection of either entry or fusion above background (**Sup.Fig.2A,B**). Supportive of our technical approach, replacing the human ACE2 signal peptide with that found in pDISPLAY had no effect on pseudotype entry or cell-cell fusion (**Sup.Fig.2**). In addition, SARS-CoV-2 entry was shown only with human ACE2, but not aminopeptidase N (APN) or dipeptidyl peptidase 4 (DPP4), the β-coronavirus group I and MERS-CoV receptors, respectively (**Sup.Fig.2**), indicating high specificity for both assays. Using the classical pseudotype approach, which models particle engagement with receptors on the surface of target cells, we demonstrated that SARS-CoV-2 Spike has a relatively broad tropism for mammalian ACE2 receptors. Indeed, we observed that pangolin, dog, cat, horse, sheep and water buffalo all sustained higher levels of entry than was seen with an equivalent human ACE2 construct (**Fig.2A**; left heatmap, first column). In contrast, all three bat ACE2 proteins we analysed (fruit bat, little brown bat and horseshoe bat) sustained lower levels of fusion than was seen with human ACE2, as did turkey and chicken ACE2, the only non-mammalian proteins tested. In accordance with previously published data on SARS-CoV and SARS-CoV-2 usage of rodent ACE2 [1, 15], rat ACE2 did not efficiently support SARS-CoV-2 particle entry. However, we observed that the ACE2 from hamsters did support pseudoparticle entry, albeit less efficiently than human ACE2.

**Figure 2:**
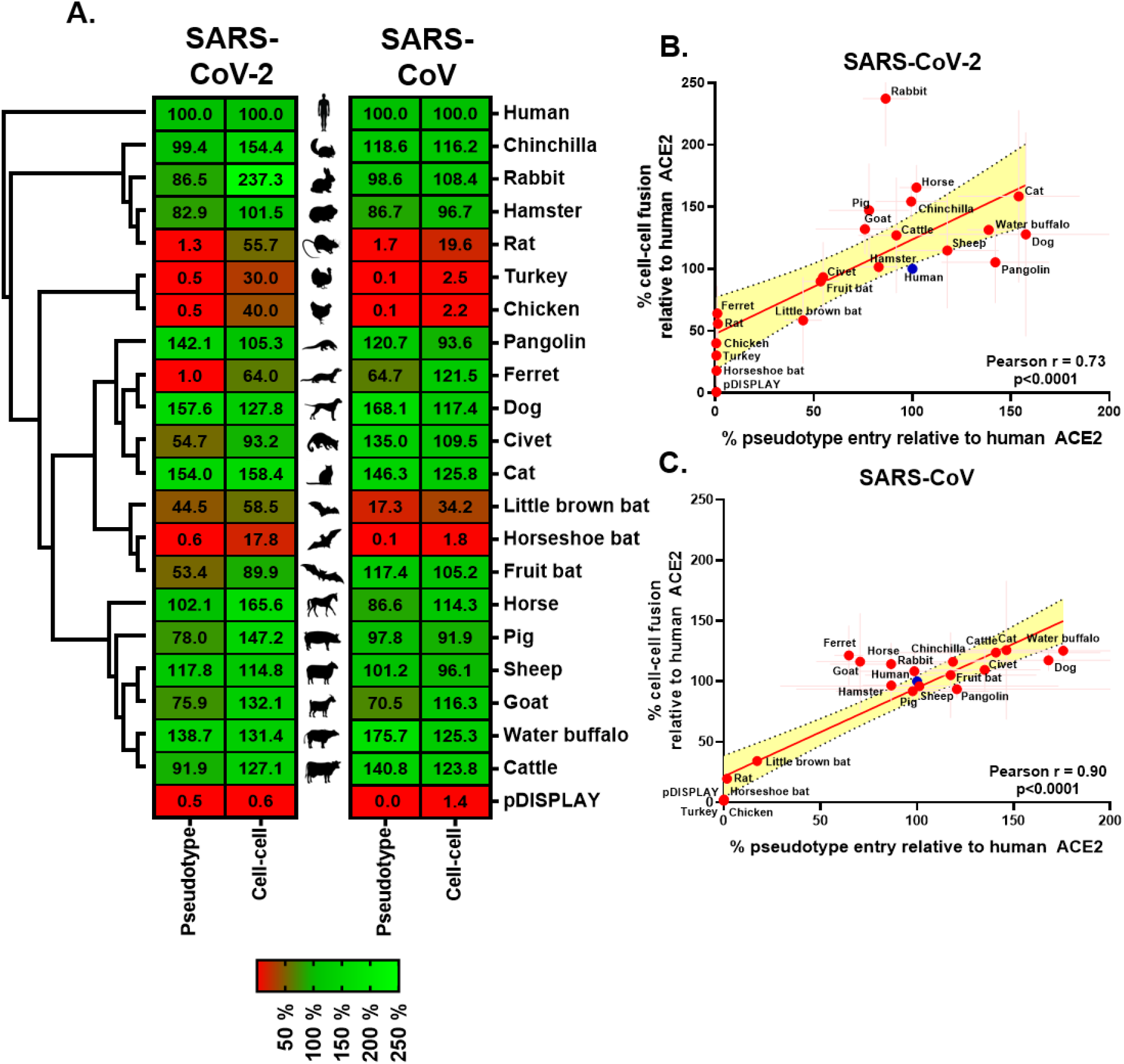
Receptor screening using surrogate entry assays identifies SARS-CoV-2 Spike as a pan-tropic viral attachment protein. **(A)** A heatmap illustrating the receptor usage profile of SARS-CoV-2 and SARS-CoV in pseudotype entry and cell-cell fusion assays with various mammalian and bird ACE2s. The data in each row is normalised to the signal seen for human ACE2 (top), with results representing the mean percentage calculated from three separate experiments performed on different days. A vector only control (pDISPLAY) was added to demonstrate specificity. Mammalian and bird ACE2s are organised, top-to-bottom based on their phylogenetic relationship (rectangular cladogram, left). **(B/C)** For both SARS-CoV-2 and SARS-CoV the respective cell-cell and pseudotype assay percentages for each ACE2 protein (relative to human ACE2) were plotted on an XY scatter graph, the Pearson correlation calculated, and a linear line of regression fitted together with 95% confidence intervals. The x and y error bars denote the standard deviation from the three experimental repeats performed on separate days.

In the separate cell-cell fusion assay, which provides both luminescence and fluorescence-based monitoring of syncytia formation, a similar trend was observed with expression of chinchilla, rabbit, hamster, pangolin, dog, cat, horse, pig, sheep, goat, water buffalo and cattle ACE2 proteins on target cells all yielding higher signals than target cells expressing human ACE2 (**Fig.2A**; left heatmap, second column). Similar to the pseudotype assay, expression of all three bat ACE2 proteins resulted in less cell-cell fusion than that seen with human ACE2. Example micrographs of GFP-positive SARS-CoV-2 Spike-induced syncytia are provided in **Sup.Fig.3**. The heatmaps presented in **Fig.2A** represent the average results from three independent pseudotype and cell-cell assay receptor usage screens (with representative data sets shown in **Sup.Fig.4**).

Combining the results from all six screens demonstrates a significant degree of concordance between the two experimental approaches. The only marked outlier is rabbit ACE2, which repeatedly generated higher signals relative to human ACE2 in the cell-cell fusion assay (**Fig.2B**). Although the high correlation (Pearson r=0.73) was unsurprising, given that both approaches rely on the same Spike-ACE2 engagement, fusogen activation and membrane fusion process (albeit at virus-cell or cell-cell interfaces), there were some marked differences in sensitivity. For the pseudotype system there was little appreciable evidence for particle entry above background levels with ferret, rat, chicken, turkey or horseshoe bat ACE2, either in vector control (pDISPLAY) transfected cells (**Fig.2A**; bottom row) or in ACE2-transfected cells infected with a ‘no glycoprotein’ pseudoparticle control, NE (**Sup.Fig.4**). However, in the cell-cell system all of these receptors permitted Spike-mediated fusion, above the background levels seen in pDISPLAY transfected cells (**Fig.2A**) or in effector cells not expressing SARS-CoV-2 Spike (**Sup.Fig.4**; No Spike), albeit at levels significantly lower than that seen for human ACE2. This suggests that these receptors, whose structures are clearly not optimal for SARS-CoV-2 entry, are still bound by the Spike protein.

To facilitate comparison with existing data for SARS-CoV, we performed all the above experiments side-by-side with SARS-CoV pseudotype and cell-cell assays (**Fig.2A,** right heatmap and **Sup.Fig.2,4**). While the receptor usage profile of SARS-CoV correlates significantly with SARS-CoV-2, both in terms of pseudotype entry (**Sup.Fig.5A**; r=0.86) and cell-cell fusion (**Sup.Fig.5B**; r=0.78), there were interesting divergences. In general, for SARS-CoV there was a better correlation between pseudotype entry and cell-cell fusion (**Fig.2B,C**; Pearson r=0.73 [SARS-CoV-2] versus r=0.90 [SARS-CoV]), with no obvious outliers and less variation between the two assays when examining receptors with low levels of associated fusion, e.g. horseshoe bat ACE2 (**Fig.2A**). These differences may be due to the differing levels of fusion seen with both viruses as well as the methodological approach taken. In our experiments SARS-CoV-2 Spike is demonstrably more fusogenic than SARS-CoV, possibly due to the presence of a furin-cleavage site between S1 and S2 [16]. Alongside a similar restriction for bird and bat ACE2 proteins, our side-by-side comparison also identified instances of varying restriction, specifically ferret, fruit bat and civet ACE2 which appear to be preferentially used by SARS-CoV (**Fig.2A** and **Sup.Fig.5B**). In summary, using two distinct technical approaches that monitor Spike-mediated receptor usage in a biologically relevant context we provide evidence that SARS-CoV-2 has a broad tropism for mammalian ACE2s. These assays demonstrate correlation between ACE2 protein sequence and fusion by SARS-CoV or SARS-CoV-2 Spike protein, plus evidence of a low affinity of SARS-CoV Spike proteins for bird or rat ACE2 and varying levels of bat ACE2 utilisation.

### A cognate ACE2 receptor is required for SARS-CoV-2 infection

High throughput and robust, surrogate assays for SARS-CoV-2 viral entry only serve to model this process and can never completely replace live virus experiments. To this end, and in order to examine the permissiveness of non-human cell lines in our cell culture collection (**Sup.Table.1**) to SARS-CoV-2, we experimentally infected a range of animal cells including those established from birds, canids, rodents, ruminants and primates with SARS-CoV-2 isolated from a patient in the UK (SARS-CoV-2 England-2/2020). Infection at a low MOI (0.001) failed to generate infectious virus in any of the cells tested, apart from two monkey cell lines (Vero E6 and Marc 145), in line with primate cells being used widely to propagate SARS-CoV-2 [17] (**Fig.3A**). Repeat infections at a higher MOI (1) in a subset of these cells (PK15, RK13, DF-1 and BHK-21) established evidence for a very low level of virus production only in the porcine cell line PK15 (**Fig.3B**). Subsequent qPCR analysis of ACE2 mRNA levels in the whole panel of cell lines, assayed using a novel panel of species-specific ACE2 primers (**Sup.Table.4**), identified only two cell lines (Vero E6 and Marc 145) with Ct values less than 25, providing a strong correlative link between ACE2 receptor expression and successful virus infection.

**Figure 3:**
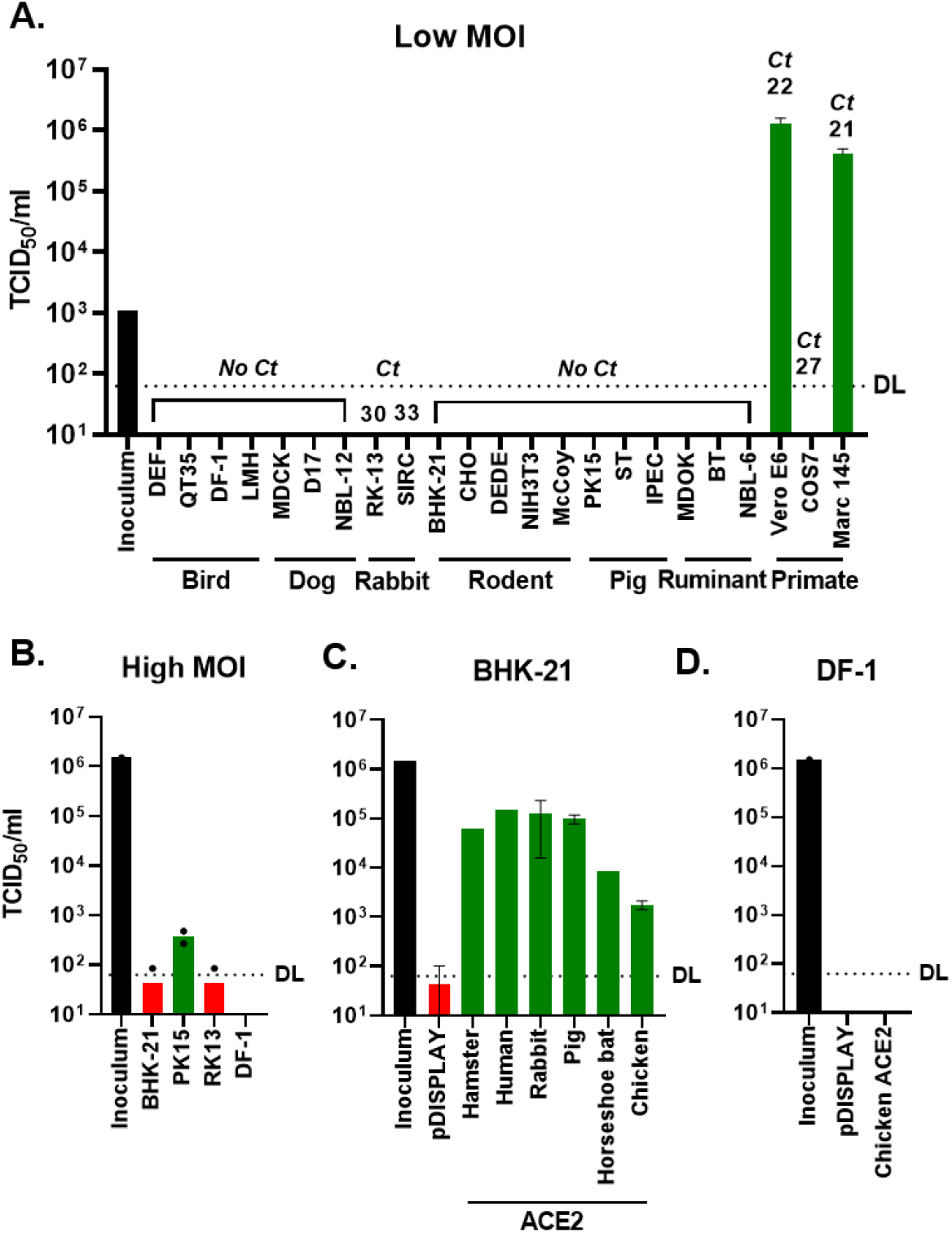
A cognate ACE2 receptor is required for SARS-CoV-2 infection. **(A)** Various cell lines derived from birds, dogs, rabbits, rodents, pigs, ruminants and primates were experimentally infected with SARS-CoV-2 at a MOI of 0.001. At 72 hours post infection the supernatants from cells were harvested and titred by TCID-50. For each cell-line RNA from uninfected cells was also extracted and RT-qPCR was performed to detect ACE2 mRNA, with the value above each line indicating the cycle when PCR positivity was achieved (Ct; cycle threshold). **(B)** Four of the same cell lines were infected again, this time at high MOI (1). **(C)** BHK-21 hamster cells were transiently transfected with ACE2 expression constructs (or a vector control [pDISPLAY]) before being infected with SARS-CoV-2 at high MOI (1). **(D)** Similarly, DF-1 cells were transfected with a chicken ACE2 expression construct or a vector control (pDISPLAY) and infected at high MOI (1). For all high MOI experiments supernatant samples were harvested at 48 hpi for titration by TCID-50. The detection limit for the TCID-50 (DL) is indicated. In all experiments the initial inoculum used for infection was titred and infections were performed in duplicate, with error bars denoting standard deviation from the mean.

We next sought to correlate the receptor usage results from our surrogate entry assays (**Fig.2**) with live virus infections. The hamster kidney cell line BHK-21, which we established as refractory to SARS-CoV-2 infection (**Fig.3A,B**), was transfected with vector alone (pDISPLAY) or a restricted panel of ACE2 constructs (hamster, human, horseshoe bat, rabbit, pig and chicken) representing the spectrum of receptor usage (**Fig.2A**). Concurrent to the infections, the expression of ACE2 in equivalently transfected cells was confirmed by western blot, flow cytometry and SARS-CoV-2 pseudotype infections (**Sup.Fig.6A-D** and **Sup.Fig.1**). Of note, for the live virus infections the high MOI (1) inoculum was removed after 1 hour with the cells thoroughly washed prior to incubation at 37 °C. Accordingly, in the BHK-21 cells transfected with carrier plasmid we saw very little evidence for virus infection and/or virus production, confirming these cells do not natively support SARS-CoV-2 infection (**Fig.3C**). For the receptors where we had previously seen high levels of cell-cell fusion (hamster, pig and rabbit) we observed robust viral replication (**Fig.3C**). Surprisingly, the two receptors included because of their ‘poor’ usage by SARS-CoV-2 Spike (horseshoe bat and chicken ACE2, **Fig.2A**) were still able to support viral replication, albeit to a lower level. Of note, regardless of the ACE2 species expressed we saw very little evidence of cytopathic effect in the infected BHK-21 cells (**Sup.Fig.6E**), despite the release of infectious virus into the supernatant (**Sup.Fig.6F**). Lastly, focusing on the unexpected observation that chicken ACE2 permitted SARS-CoV-2 entry into cells, we investigated whether chicken DF-1 cells over-expressing ACE2 could support viral replication. Whilst western blot and flow cytometry demonstrated successful ACE2 over-expression (**Sup.Fig.6B,D**) we did not see any evidence of viral replication in these cells, either because of inefficient chicken ACE2 receptor usage or a post-entry block to SARS-CoV-2 replication (**Fig.3D**). In summary, SARS-CoV-2 is able to use a range of non-human ACE2 receptors to enter cells. Furthermore, when a cognate ACE2 is provided the virus can replicate efficiently in the normally refractory hamster cell line BHK-21.

## Discussion

Recognising animals at risk of infection and/or identifying the original or intermediate hosts responsible for the SARS-CoV-2 pandemic are important goals for ongoing COVID-19 research. In addition, there is a requirement to develop appropriate animal models for infection that, if possible, recapitulate the hallmarks of disease seen in people. Importantly, high-resolution structures of human ACE2 in complex with the Spike RBD [5, 6, 10, 11] can help us to understand the genetic determinants of SARS-CoV-2 host-range and pathogenesis. In particular, differences in receptor usage between closely related species provides an opportunity to pinpoint amino acid substitutions at the interaction interface that inhibit Spike protein binding and thus fusion.

One example of closely related ACE2 sequences differing in their utilisation by SARS-CoV-2 Spike comes from the comparison of rat and hamster ACE2. Although a number of animal models have been investigated for SARS-CoV-2, including non-human primates, ferrets and cats [18, 19], the use of small animals, in particular rodents, has proved more challenging as murine and rat ACE2 support lower levels of β-coronavirus entry [1, 15]. For SARS-CoV this problem was circumvented with the development of transgenic mice expressing human ACE2 [20] or mouse-adapted SARS-CoV [21, 22]. Consistent with previously published data on SARS-CoV rodent ACE2 interactions, we showed that rat ACE2 does not support SARS-CoV-2 mediated fusion (**Fig.2A**). However, our finding that hamster ACE2 allows entry of SARS-CoV-2 (**Fig.2A**) indicates this animal is a suitable model for infection, consistent with recent *in vivo* studies demonstrating experimental infection of these animals [23]. Comparison of the hamster and rat sequences (**Fig.4A**) identified multiple substitutions at the RBD interaction interface that might explain this variable receptor tropism (listed as hamster to rat): Q24K, T27S, D30N, L79I, Y83F, K353H. Except for L79I, which is similarly substituted in pangolin and pig ACE2, all of these substitutions are likely to reduce Spike RBD binding. Q24K and Y83F substitutions would both result in the loss of hydrogen bonds with the side chain of SARS-CoV-2 RBD residue N487 (**Fig.4B**). Residue D30 is acidic in all ACE2 proteins that are efficiently utilised by SARS-CoV-2 Spike, and its substitution to asparagine would remove the salt bridge formed with K417 of the RBD. Lastly, the T27S substitution would remove the threonine side chain methyl group that sits in a hydrophobic pocket formed by the side chains of RBD residues F456, Y473, A475 and Y489. Thus, multiple substitutions are predicted to inhibit Spike binding to rat ACE2 when compared with the closely related hamster protein. Of note, the hamster cell lines we used in our study (BHK-21 and CHO) are likely refractory to infection simply because they express low levels of ACE2 mRNA (**Fig.3A**, qPCR data). Interestingly, the high level of cell-cell fusion seen with rabbit ACE2 indicates that lagomorphs may also represent a good model organism for SARS-CoV-2 pathogenesis.

A second example of different receptor usage between closely related species can be seen with bat ACE2 (**Fig.2A, 4A**). The apparent lack of tropism for bat ACE2 proteins we observed was surprising as there is previous evidence of SARS-CoV-2 infection of bat ACE2 expressing cells *in vitro* [1] and *in vitro* binding experiments suggest that the SARS-CoV-2 RBD binds bat ACE2 with high affinity [24]. Since the exact origin of SARS-CoV-2 is currently unknown, but widely accepted to be a Chiroptera species, we included ACE2 proteins from a broad range of bats in our study. While none support SARS-CoV-2 fusion to the same levels as humans, there are dramatic differences in the ability of SARS-CoV-2 Spike to utilise ACE2 from horseshoe bats versus fruit bats and little brown bats (**Fig.2A**). As discussed earlier, the closest known relative of SARS-CoV-2, RaTG13, was isolated from intermediate horseshoe bat (*Rhinolophus affinis*). Unfortunately, the ACE2 sequence from this species was not available for use in our study; however, we did include an ACE2 from the closely related least horseshoe bat (*Rhinolophus pusillus*). Although this protein supported the lowest levels of fusion of any bat ACE2 tested in our study, it still supported a low level of SARS-CoV-2 replication with live virus (**Fig.3C**). As in rat ACE2, horseshoe bat and fruit bat ACE2 have a lysine residue at position 24 that would disrupt hydrogen bonding to N487 of the SARS-CoV-2 RBD and introduce a charge (**Fig.4A,B**). Little brown bats have the hydrophobic residue leucine at this position, which could not form the hydrogen bond to N487 but which is present in ACE2 from several species that support high levels of fusion, suggesting that loss of the hydrogen bond is less deleterious to Spike protein binding than introduction of the lysine positive charge. Fruit bats conserve a T27 whereas little brown bats have the bulkier isoleucine residue and horseshoe bats have a bulky charged lysine residue in this position, both of which are likely to clash with the F456-Y473-A475-Y489 hydrophobic pocket of the RBD, with the lysine substitution likely to be more deleterious due to the introduction of the positive charge. Like rats, horseshoe bat N30 would be unable to form a salt bridge with RBD K417. Substitution of Q42 with glutamate in little brown bat may be detrimental to Spike binding as it would disrupt the hydrogen bond to the backbone carbonyl oxygen of RBD residue G446. The other substitutions between bat ACE2 proteins and other mammals are likely to be benign. Little brown bats, horseshoe bats, pangolins and horses all share a serine as ACE2 residue 34, suggesting that serine in this position does not abolish Spike binding, and it is likely that the threonine at this position (fruit bat ACE2) would likewise be tolerated. Similarly, the Y41H substitution present in little brown bat ACE2 is also present in horse ACE2, suggesting that it does not prevent binding. Therefore, all bat ACE2 proteins have substitutions that impair SARS-CoV-2 Spike binding to different degrees, but it seems likely that the E30N substitution (shared only by rat ACE2) is the most likely cause of the severely impaired binding of SARS-CoV-2 Spike to horseshoe bat ACE2.

**Figure 4:**
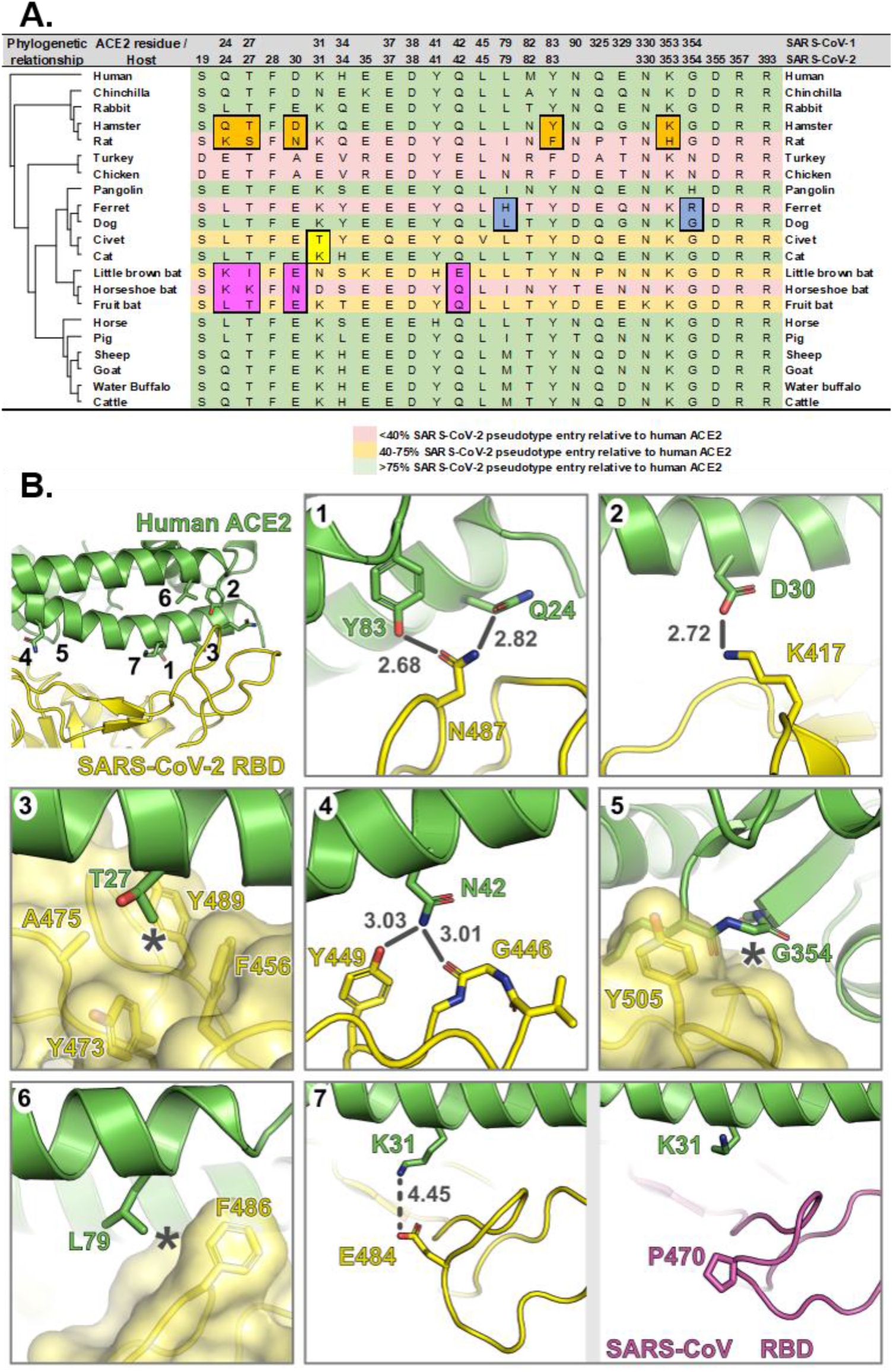
Substitutions at the interface between SARS-CoV-2 RBD and mammalian ACE2 proteins impact receptor utilisation. **(A)** Residues of mammalian ACE2 sequences used in this study that are predicted to interact with the RBD SARS-CoV and −2, based on the structures of human ACE2 in complex with SARS-CoV [28] and SARS-CoV-2 [6]. Differences between closely-related species that may impact RBD binding are highlighted. **(B)** Interface between human ACE2 (green) and SARS-CoV-2 RBD (yellow). Insets 1 to 7 show molecular interactions discussed in the main text. Bonds that may be disrupted are shown as grey lines, with bond distances in grey text, and hydrophobic interactions that may be disrupted are marked with asterisks. The right and left hand panels of inset 7 show human ACE2 in complex with (left) SARS-CoV-2 [10] or (right) SARS-CoV RBD [28].

Interestingly, a similarly ‘poor’ tropism for bat ACE2 was also reported for SARS-CoV following its emergence in 2002 [25]. Specifically, coronaviruses closely related to SARS-CoV that were isolated directly from bats were shown to not efficiently use either human or civet ACE2 [25].

This is consistent with large shifts in receptor usage occurring during coronavirus species jumps, either directly into humans or more likely via intermediate reservoirs. During the SARS-CoV epidemic, where civets were identified as the intermediate reservoir of infection, a shifting pattern of increasing and decreasing ACE2 usage was observed in individual isolates of SARS-CoV taken from civets and humans (although they shared ~99% similarity to each other), providing evidence for adaptation to individual host receptors [12, 26] with a particular focus on differential adaptation to human ACE2 residues K31 (T31 in civets) and K353. Interestingly, correlation analysis of SARS-CoV and SARS-CoV-2 pseudotype entry highlighted civet ACE2 as being strongly favoured by SARS-CoV, perhaps a legacy of this period of adaptation in an intermediate host (**Sup.Fig.4A**). Although data analysis of this type between related viruses might represent a mechanism for identifying intermediate reservoirs, similar outliers that favoured SARS-CoV-2 entry were not evident in our study. Unfortunately, the lack of similarly closely related SARS-CoV-2 isolates from this outbreak’s origin in Hubei makes detailed interpretation of this virus’s adaptation to human ACE2 difficult at this time. Recently, pangolins were demonstrated to harbour SARS-related coronaviruses, implicating these animals as the potential intermediate [27]. Although this hypothesis is supported by our receptor usage data, these isolates are probably too dissimilar from SARS-CoV-2 (90% at the genomic level) to have been the immediate source of the current pandemic. Based on our findings (**Fig.2A**) it is theoretically possible that several different animals could have acted as the intermediate host for this virus. However, without original isolates from Hubei it may not be possible to easily identify the animal or animals that seeded the original epidemic.

A third example of ACE2 usage by SARS-CoV-2 differing dramatically between closely related species is dog and ferret. It was surprising that entry of SARS-CoV-2 pseudotypes into cells was heavily restricted by ferret ACE2 (c. 1% of human levels, **Fig.2A**), despite this animal developing established signs of infection following experimental challenge [19] and its clear potential for use as an animal model [18]. Substitution of G354 in dog ACE2 with the bulky, charged residue arginine in ferret is likely to decrease binding efficiency, although we note that pangolin ACE2 has a His residue in this position and retains binding to Spike (**Fig.4A,B**). Similarly, substitution of L79 in dog ACE2 with histidine in ferret would be likely to disrupt hydrophobic interactions with the side chain of Spike F486. Comparison of mammalian ACE2 receptors usage by SARS-CoV versus SARS-CoV-2 can also be coupled to inspection of the available ACE2:RBD co-structures [5, 6, 10, 11, 28] to obtain further molecular insights into Spike binding. The long arginine side chain of SARS-CoV residue 426, which is the structural equivalent to SARS-CoV-2 N439, makes a salt bridge with E329 of human ACE2 and interacts with the side chain of ACE2 Gln325. This results in a larger binding footprint for SARS-CoV on human ACE2 when compared to SARS-CoV-2 (see, for example, Figure 3 in [6]). It is therefore striking that ferret ACE2 is not used efficiently by SARS-CoV-2 for fusion, while ferret ACE2 can support SARS-CoV-mediated fusion. The enhanced usage may arise from a salt bridge being formed between ferret ACE2 residue E325 and SARS-CoV R426, which is not possible in SARS-CoV-2 where the equivalent residue is asparagine. This additional salt bridge may therefore ‘rescue’ some of the binding loss caused by the deleterious substitutions listed above. For ferret ACE2 the dichotomy between established *in vivo* infection but poor receptor usage *in vitro* is perplexing. However, it is consistent with our observation that chicken ACE2 can support SARS-CoV-2 infection at high MOI (**Fig.3C**) and suggests that low levels of SARS-CoV-2 Spike-mediated fusion are sufficient for cell entry. The widescale availability of ferrets as an animal model of infection could represent an excellent opportunity to study in-host adaptation of SARS-CoV-2 to the ACE2 receptor, mimicking the steps that lead up to the establishment of intermediate reservoirs in the wild.

While not as dramatic as for the species listed above, differences in SARS-CoV-2 receptor utilisation are also seen for cat versus civet ACE2. The most likely candidate causative substitution in civet ACE2 is K31T, where a complementary long-range charge interaction with RBD E484 is lost. In SARS-CoV the entire loop between residues 460–472 (equivalent to SARS-CoV-2 residues 473–486) is reordered and there is not a glutamate or aspartate residue in this local vicinity. We would therefore not predict cat ACE2 to bind SARS-CoV Spike more strongly than civet ACE2, consistent with our fusion assay data.

In the process of finalising this manuscript two papers were released as preprints, also examining the receptor usage of various non-human ACE2s with surrogate virus entry assays (lentiviral pseudotypes) [29, 30]. While these studies did not perform corresponding examination of cell-cell fusion or follow up experiments with SARS-CoV-2 live virus there is a strong correlation between their findings and ours, namely the broad tropism of SARS-CoV-2 Spike. Notably, all three research data sets concur that human and several non-human ACE2 proteins support similar levels of utilisation by SARS-CoV-2 Spike, in contrast to a recent report that claimed preferential binding to the human ACE2 protein [31]. One interesting conclusion, drawn specifically from our technical approach, is that pseudotype assays alone may not fully capture the receptor tropism of SARS-CoV-2. Like Li et al. [29], we observed very little evidence of SARS-CoV-2 pseudoparticle entry into cells expressing chicken ACE2; nevertheless, live virus was able to enter and replicate in equivalent cells, albeit to a lesser extent than with human ACE2. There are various interpretations for this result, including roles for additional host factors involved in entry such as the recently identified SARS-CoV-2 Spike-neurophilin-1 interaction [32]. Another explanation is that SARS-CoV-2 Spike is not correctly presented on the surface of HIV pseudotypes (either numerically or spatially) and this therefore might not confer full wild-type-virus-like infectivity to lentiviral pseudoparticles. This could be especially important for coronaviruses which contain other envelope viral proteins E and M, which spontaneously form VLPs when expressed with Spike. Unfortunately, while VLPs may more accurately reflect the number and conformation of viral proteins in the live virus particle they cannot easily be manipulated to encode a reporter gene, such as Firefly luciferase. As such our cell-cell system or indeed live virus may therefore be more appropriate for probing low affinity interactions between atypical host ACE2s and coronavirus Spike proteins. While more evidence is required to examine the significance of cell-cell spread (syncytia formation) of SARS-CoV-2 *in vitro* and *in vivo*, the quantitative assay we have developed promises to be a robust tool for supporting these efforts. Similarly, examining whether the polybasic cleavage site found in SARS-CoV-2 Spike provides a selective advantage to this virus, e.g. by allowing enhanced spread of the virus through cell-cell fusion, is the source of ongoing investigations in our laboratory. Similar trends have been observed for related murine and bovine coronaviruses [33–36] and our quantitative cell-cell assay may also prove useful in characterising this aspect of the viral life cycle.

Using both live virus and quantitative assays that model Spike-receptor usage we have demonstrated that SARS-CoV-2 has a broad tropism for mammalian ACE2s. These findings are supported by results from experimental challenge infections, including cats that are susceptible to infection and chickens that are not [19], as well as evidence of community-based reverse zoonotic infections observed in cats and dogs [37, 38]. At the time of writing, the epidemiological significance of these infections remains to be determined, for example whether they represent one-off spill-over events without onward transmission, or alternatively evidence for the existence of animal reservoirs. The latter scenario would have significant epidemiological implications to human populations recovering from the first wave of the SARS-CoV-2 pandemic. Interestingly, certain animals where we demonstrated efficient receptor usage, e.g. pigs and dogs (**Fig.2A**), appear less susceptible to experimental challenge [19]. It should be noted that particle entry represents only the first step in zoonotic spill-over; indeed, multiple virus-host interactions contribute to define virus host-range and pathogenesis. It may be that the intra-cellular environment of specific hosts cannot sustain SARS-CoV-2 infection, either through the absence of an important virus-host interaction or the presence of effective mechanisms of innate immune restriction. In this case SARS-CoV-2 may enter cells efficiently but fail to replicate to significant levels to support onward transmission, bring about clinical signs or induce immunopathological sequalae. In conclusion, SARS-CoV-2 exhibits a broad tropism for mammalian ACE2s. More thorough investigation, including heightened virus surveillance and detailed experimental challenge studies, are required to ascertain whether livestock and companion animals could act as reservoirs for this disease or as targets for reverse zoonosis.

## Methods

### Cell lines

Cell lines representing a broad range of animal species were used to determine the host range/tropism of SARS-CoV-2 (**Sup.Table.1)** (Cell Culture Central Services Unit, The Pirbright Institute). Cells were maintained in complete medium supplemented with either 10% horse or BVDV/FMDV-negative foetal bovine serum (FBS) (Gibco), 1% non-essential amino acids, 1mM sodium pyruvate solution (Sigma-Aldrich), 2mM L-Glutamine (Sigma-Aldrich) and 1% Penicillin/Streptomycin, 10,000U/mL (Life Technologies Ltd). Additional supplements and cell culture medium for each cell line are summarised in **Sup.Table.1.** All cells were incubated at 37°C in a humidified atmosphere of 5% CO_2_.

#### Cells used for entry studies or fusion assays

HEK293T cells stably expressing a split *Renilla* luciferase-GFP plasmid (rLuc-GFP 1-7) and BHK-21 cells were maintained in DMEM-10%: Dulbecco’s Modified Eagle’s Medium, DMEM (Sigma-Aldrich) supplemented with 10% FBS (Life Science Production), 1% 100mM sodium pyruvate (Sigma-Aldrich), 1% 200mM L-Glutamine (Sigma-Aldrich), and 1% Penicillin/Streptomycin, 10,000U/mL (Life Technologies Ltd). Stable cell lines were generated, as described previously, using a lentiviral transduction system under 1μg/mL puromycin (Gibco) selection (Thakur 2020, manuscript in preparation and [39]).

### Viruses and virus titre quantification

SARS-CoV-2 England-2/2020 was isolated from a patient in the UK and a passage 1 stock was grown and titred in Vero E6 cells by PHE (kindly provided to The Pirbright Institute by Prof. Miles Carroll). A master stock of virus was passaged to P2 in Vero E6 at a MOI of 0.001 in DMEM/2% FBS and used for all virus assays, following a freeze-thaw cycle at −80°C. Stocks were titred by plaque assay on Vero E6 cells using a 1xMEM/0.8% Avicel/2% FBS overlay, fixed using formaldehyde and stained using 0.1% Toluidine Blue. All infections were performed in ACDP HG3 facilities by trained personnel.

### Plasmids

Codon optimised ACE2-expressing plasmids from a range of animal species were synthesised and cloned into pDISPLAY (BioBasic) (**Sup.Table.2)**. A codon optimised SARS-CoV protein sequence was synthesised and cloned into pcDNA3.1+ (BioBasic) while the pCAGGS plasmid expressing codon optimised SARS-CoV-2 was obtained from NIBSC, UK (**Sup.Table.3)**.

### Infections

#### Initial screen

Cells listed in **Sup.Table.1** were seeded at a density of 1×10^5^ cells/well in a 24-well plate (Nunc) and 24h later infected with SARS-CoV-2 England-2/2020. Briefly, media was removed and the cells washed once with complete DMEM supplemented with 2% FBS. Cells were then infected at MOI 0.001 and incubated at 37°C for 1h. Following this, inoculum was removed, cells washed twice with PBS, complemented with cell maintenance media and incubated for 72h at 37°C. Supernatant was collected at 72 h post infection and frozen at −80°C until required. Cells were fixed with formaldehyde for 30 mins and then stained with 0.1% Toluidine Blue (Sigma-Aldrich).

#### Receptor usage screen

BHK-21 and DF-1 cells were plated at 1×10^5^ cells/well in 24-well plates (Nunc). The following day, cells were transfected with 500ng of a subset of ACE2 expression constructs (human, hamster, rabbit, pig, chicken, horseshoe bat) or mock transfected with an empty vector (pDISPLAY) in OptiMEM (Gibco) using TransIT-X2 transfection reagent (Mirus) according to the manufacturer’s recommendations. Following this, cells were infected at MOI 1 as described above and supernatants collected at 72h post infection and frozen at −80°C until required.

### SARS-CoV-2 and SARS-CoV pseudoparticles infections

#### Pseudoparticle generation

Lentiviral based pseudoparticles were generated in HEK293T producer cells, seeded in 6-well plates at 7.5×10^5^/well one day prior to transfecting with the following plasmids: 600ng p8.91 (encoding for HIV-1 gag-pol), 600ng CSFLW (lentivirus backbone expressing a firefly luciferase reporter gene) and either 25ng of SARS-CoV-2 Spike or 500ng SARS-CoV in OptiMEM (Gibco) (**Sup.Table.3)** with 10μL PEI, 1μg/mL (Sigma) transfection reagent. No glycoprotein controls (NE) were also set up using empty plasmid vectors (25ng pCAGGS for SARS-CoV-2 and 500ng pcDNA3.1 for SARS-CoV) and all transfected cells were incubated at 37°C, 5% CO_2_. The following day, the transfection mix was replaced with DMEM-10% and pooled harvests of supernatants containing SARS-CoV-2 pseudoparticles (SARS-CoV-2 pps) and SARS-CoV pseudoparticles (SARS-CoV pps) were taken at 48 and 72h post transfection, centrifuged at 1,300 x *g* for 10 mins at 4°C to remove cellular debris, aliquoted and stored at −80°C. HEK293T target cells transfected with 500ng of a human ACE2 expression plasmid (Addgene) were seeded at 2×10^4^ in 100μL DMEM-10% in a white-bottomed 96-well plate (Corning) one day prior to infection. SARS-CoV-2 pp and SARS-CoV pp along with their respective NE controls were titrated 10-fold on target cells and incubated for 72h at 37°C, 5% CO_2_. To quantify firefly luciferase, media was replaced with 50μL Bright-Glo™ substrate (Promega) diluted 1:2 with serum-free, phenol red-free DMEM, incubated in the dark for 2 mins and read on a Glomax Multi+ Detection System (Promega).

#### Receptor usage screens

BHK-21 cells were seeded in 48-well plates at 5×10^4^/well in DMEM-10% one day prior to transfection with 500ng of different species ACE2-expression constructs or empty vector (pDISPLAY) (**Sup.Table.2)** in OptiMEM and TransIT-X2 (Mirus) transfection reagent according to the manufacturer’s recommendations. The next day, cells were infected with SARS-CoV-2 pp/SARS-CoV pp equivalent to 10^6^-10^7^ relative light units (RLU), or their respective NE controls at the same dilution and incubated for 48h at 37°C, 5% CO_2_. To quantify Firefly luciferase, media was replaced with 100μL Bright-Glo™ substrate (Promega) diluted 1:2 with serum-free, phenol red-free DMEM. Cells were resuspended in the substrate and 50μL transferred to a white-bottomed plate in duplicate. The plate was incubated in the dark for 2 mins then read on a Glomax Multi+ Detection System (Promega) as above. CSV files were exported onto a USB flash drive for analysis. Biological replicates were performed three times.

### Cell-cell fusion assays

HEK293T rLuc-GFP 1-7 [40] effector cells were transfected in OptiMEM (Gibco) using Transit-X2 transfection reagent (Mirus), as per the manufacturer’s recommendations, with SARS-CoV-2 (250ng), SARS-CoV (1000ng) (**Sup.Table.3**) or mock-transfected with empty plasmid vector (pCAGGS for SARS-CoV-2 and pcDNA3.1+ for SARS-CoV). BHK-21 target cells were co-transfected with 500ng of different ACE2-expressing constructs (**Sup.Table.2**) and 250ng of rLuc-GFP 8-11 plasmid. For SARS-CoV-2 cell-cell fusion assays, target cells were also transfected with 25ng of transmembrane protease serine 2 (TMPRSS2) for 48h. SARS-CoV-2 effector cells were washed once with PBS and resuspended in phenol red-free DMEM-10%. SARS-CoV effector cells were washed twice with PBS and incubated with 3μg/ml of TPCK-treated trypsin (Sigma-Aldrich) for 30 mins at 37°C before resuspending in phenol red-free DMEM-10%. Target cells were washed once with PBS and harvested with 2mM EDTA in PBS before co-culture with effector cells at a ratio of 1:1 in white 96-well plates to a final density of 4×10^4^ cells/well in phenol red-free DMEM-10%. Quantification of cell-cell fusion was measured based on *Renilla* luciferase activity, 18h (SARS-CoV-2) or 24h (SARS-CoV) later by adding 1μM of Coelenterazine-H (Promega) at 1:400 dilution in PBS. The plate was incubated in the dark for 2 mins then read on a Glomax Multi+ Detection System (Promega) as above. CSV files were exported onto a USB flash drive for analysis. GFP fluorescence images were captured every 2h for 24h using an Incucyte S3 real-time imager (Essen Bioscience, Ann Arbor, MI, USA). Cells were maintained under cell culture conditions as described above. Assays were set up with three or more biological replicates for each condition, with each experiment performed three times.

### Western blotting

BHK-21 cells were transfected using Transit-X2 transfection reagent (Mirus), as per the manufacturer’s instructions with 500ng of different ACE2-expression constructs (**Sup.Table.2**) or mock-transfected with empty plasmid vector (pDISPLAY). All protein samples were generated using 2x Laemmli buffer (Bio-Rad) and reduced at 95°C for 5 mins 48h post-transfection. Samples were resolved on 7.5% acrylamide gels by SDS-PAGE, using semi-dry transfer onto nitrocellulose membrane. Blots were probed with mouse anti-HA primary antibody (Miltenyi Biotech) at 1:1,000 in PBS-Tween 20 (PBS-T, 0.1%) with 5% (w/v) milk powder overnight at 4°C. Blots were washed in PBS-T and incubated with anti-mouse secondary antibody conjugated with horseradish peroxidase (Cell Signalling) at 1:1,000 in PBS-T for 1h at room temperature. Membranes were exposed to Clarity Western ECL substrate (Bio-Rad Laboratories) according to the manufacturer’s guidelines and exposed to autoradiographic film.

### Flow cytometry

BHK-21 cells were transfected using Transit-X2 transfection reagent (Mirus), as per the manufacturer’s instructions with 500ng of each ACE2-expression construct (**Sup.Table.2**) or mock-transfected with empty plasmid vector (pDISPLAY) for 48h. Cells were resuspended in cold PBS and washed in cold stain buffer (PBS with 1% BSA (Sigma-Aldrich), 0.01% NaN3 and protease inhibitors (Thermo Scientific)). Cells were stained with anti-HA PE-conjugated antibody (Miltenyi Biotech) at 1:50 dilution for 1×10^6^ cells for 30 mins on ice, washed twice with stain buffer and fixed in 2% paraformaldehyde for 20 mins on ice. Fixed cells were resuspended in PBS before being analysed using the MACSQuant^®^ Analyzer 10 (Miltenyi Biotech) and the percentage of PE-positive cells was calculated by comparison with unstained and stained mock-transfected samples. Positive cells were gated as represented in **Sup.Fig.1** and the same gating strategy was applied in all experiments.

### RNA extraction and ACE2 qPCR quantification

Total cellular RNA was extracted from cell lines in **Sup.Table.1** using a QIAGEN RNeasy RNA extraction kit and mRNA was then detected with SYBR-green based qPCR, using a standard curve for quantification on a Quant studio 3 thermocycler. Luna^®^ Universal qPCR Master Mix (NEB) was used to quantify mRNA levels for each cell line. RNA was first transcribed using SuperScript II Reverse Transcriptase (Thermo Fisher), with oligo dT primers and 50ng of input RNA in each reaction. All the reactions were carried out following the manufacturer’s instructions and in technical duplicate, with the melt curves analysed for quality control purposes. Conserved cross-species ACE2 primers used for each cell line are found in **Sup.Table.4**.

### Structural analysis

Molecular images were generated with an open source build of PyMOL (Schrödinger) using the crystal structure of SARS-CoV-2 RBD in complex with human ACE2 (PDB ID 6M0J) [10] that had been further refined by Dr Tristan Croll, University of Cambridge (https://twitter.com/CrollTristan/status/1240617555510919168). To analyse the conservation of mammalian ACE2 receptor sequences, representative sequences were identified via a PSI-BLAST [41] search of the UniRef90 database [42] and filtering for the class mammalia (taxid:40674). The selected sequences were aligned using MAFFT [43] and evolutionary conservation of amino acids was mapped onto the ACE2 structure using ConSurf [44].

### Phylogenetic analysis

ACE2 amino acid sequences were translated from predicted mRNA sequences or protein sequences (**Sup.Table.2**). The predicted guinea pig mRNA sequence was more divergent than expected and contained a premature stop codon. For the purposes of this research, five single nucleotides were added, based on the most closely related sequence (chinchilla), to allow a full-length mature protein to be synthesised. It is not clear if the guinea pig has a functional ACE2, or if the quality of the genomic data is very low, but overall confidence in this sequence is low. The other divergent sequence was turkey as the 3’ end was not homologous with other vertebrate ACE2 receptors. This appeared to be a mis-annotation in the genome as the 3’ end showed very high identity to the collectrin gene. The missing 3’ of the gene was found in the raw genome data and assembled with the 5’ region to make a full ACE2 sequence. Twenty-three nucleotide base pairs were missing between these regions; these were taken from chicken as the most closely related sequence. Phylogenetic analysis of the final dataset was inferred using the Neighbor-Joining method [45] conducted in MEGA7 [46] with ambiguous positions removed. The tree is drawn to scale and support was provided with 500 bootstraps.

### Data handling and statistical analysis

GraphPad Prism v8.2.1 (GraphPad Software) was used for graphical and statistical analysis of data sets. Flow cytometry data was analysed using FlowJo software v10.6.2 (BD).

## Author Contributions

Conceptualisation: DB, HJM, EB; Methodology: DB, CC, NT, JAH, ID, HS, SCG; Investigation: CC, NT, SH, JK, LL, DaB, SB, PSL, AD, PH, MV, CT, JAH, HS, ID, SCG, DB; Resources: DB, ID, HS; Writing—Original Draft Preparation: DB, CC, NT, SCG; Writing—Review and Editing: All authors; Project Administration: DB, CC, NT; Funding Acquisition: DB.

## Acknowledgements

We would like to thank the following for assistance establishing the SARS-CoV pseudotype systems: Ed Wright (University of Sussex), Nigel Temperton and Simon Scott (University of Kent), Brian Willett (University of Glasgow Centre for Virus Research), Emma Bentley and Giada Mattiuzzo (National Institute for Biological Standards and Control [NIBSC]) and Michael Letko (National Institute of Allergy and Infectious Diseases). We also acknowledge the support of Nadine Lewis (BioBasic) for help with gene synthesis as well as the The Pirbright Institute’s Flow Cytometry Facility and The Pirbright Institute Cell Servicing Unit.

## Funding

This work was supported by the following grants to DB: a UK Research and Innovation (UKRI; www.ukri.org) Medical Research Council (MRC) New Investigator Research Grant (MR/P021735/1), a UKRI Biotechnology and Biological Sciences Research Council (BBSRC, www.ukri.org) project grant (BB/R019843/1) and Institute Strategic Programme Grant (ISPG) to The Pirbright Institute (BBS/E/I/00007034, BBS/E/I/00007030 and BBS/E/I/00007039) and an Innovate UK Department for Health and Social Care project (SBRI Vaccines for Global Epidemics – 795 Clinical; Contract 971555 ‘A Nipah vaccine to eliminate porcine reservoirs and safeguard human health’). SCG is a Sir Henry Dale Fellow, jointly funded by the Wellcome Trust and the Royal Society (098406/Z/12/B).

## Competing Interests Statement

The authors declare no competing interests. The funders played no role in the conceptualisation, design, data collection, analysis, decision to publish, or preparation of the manuscript.

## Materials & Correspondence

Correspondence and material requests to Dr. Dalan Bailey.

---

**Supplemental Figure 1:**
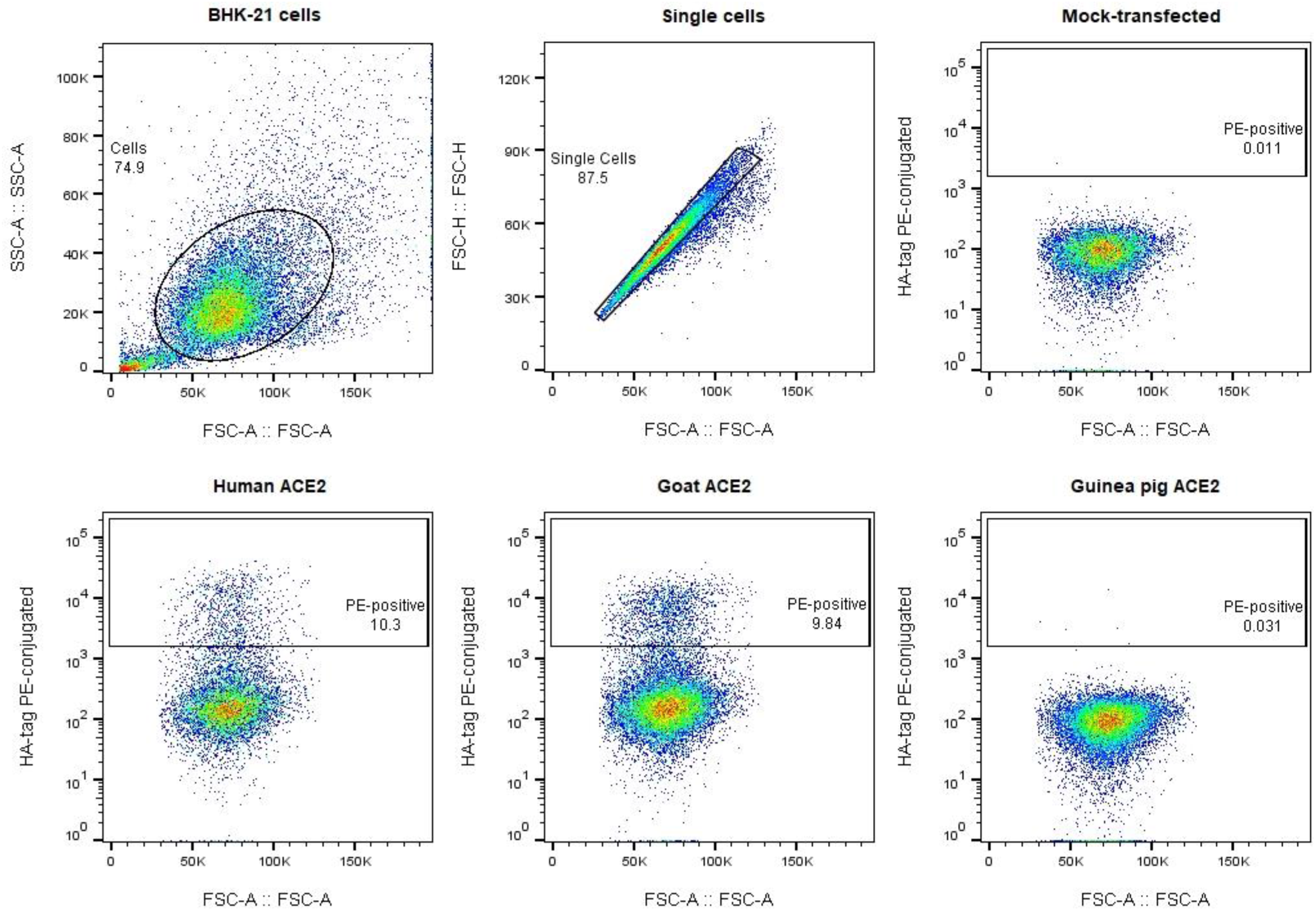
Gating strategy for flow cytometry analysis of ACE2-expressing constructs. BHK-21 cells were transfected with a panel of species-specific ACE2-expressing constructs (see **Sup.Table.2**). Cells were surface stained with anti-HA PE conjugated antibody. Live and singlet BHK-21 were gated as PE-positive, relative to mock-transfected cells (**top panel**). Representative datasets are shown for human, goat and guinea pig ACE2 surface staining (**bottom panel**).

**Supplemental Figure 2:**
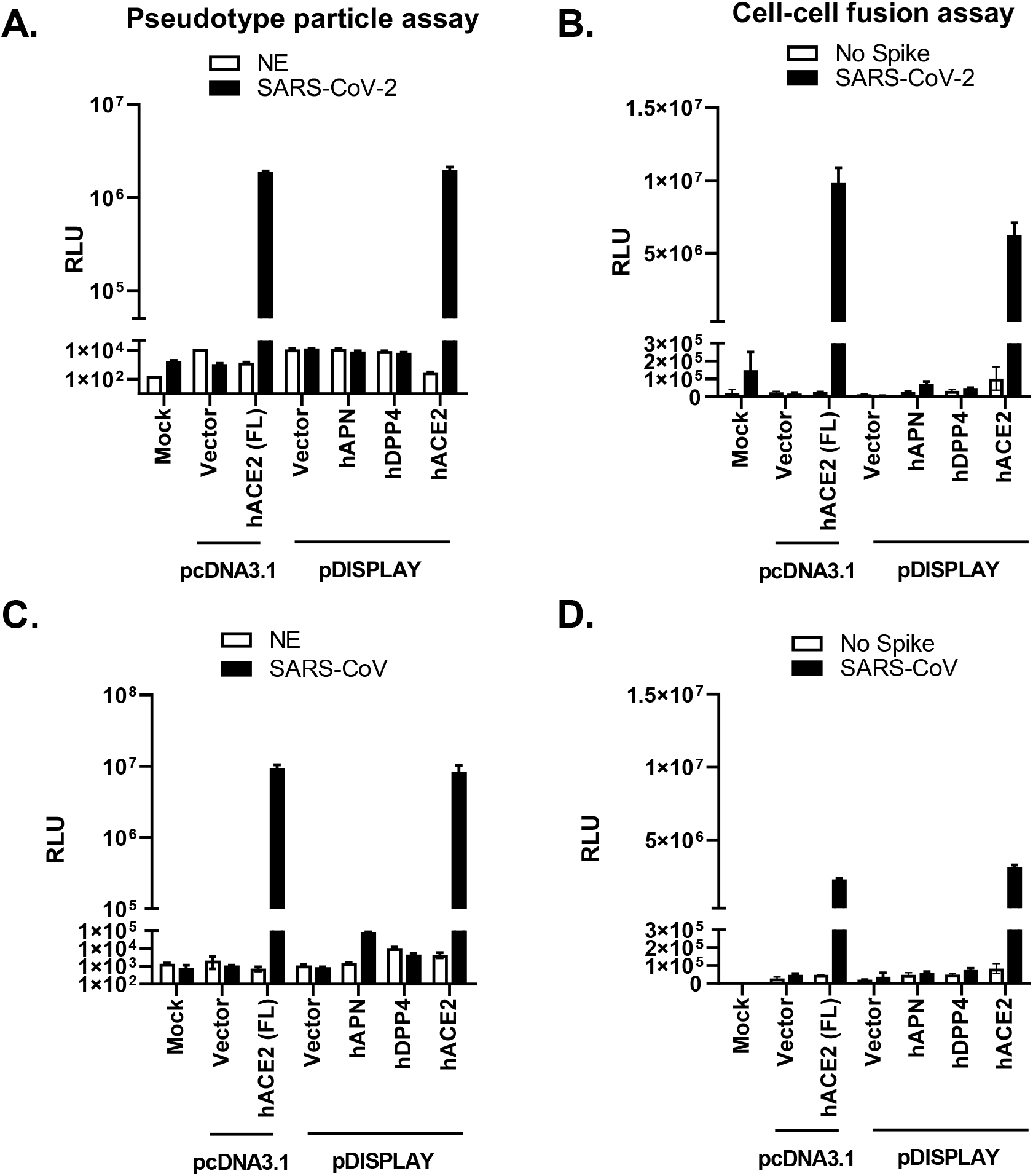
Establishment of SARS-CoV-2 and SARS-CoV entry assays. **(A-D)** Pseudotype and cell-cell fusion assays were established for SARS-CoV-2 (**A,B**) and SARS-CoV (**C,D**) using multiple internal controls. For the pseudotype assays non-enveloped (NE) lentiviral particles were generated, i.e. vector plasmid in place of a viral glycoprotein, to examine background levels of pseudoparticle entry. For the cell-cell fusion assay mock-transfected effector cells were used (No Spike) to examine background levels of cell-cell fusion. In all subsequent experiments ‘NE’ and ‘No Spike’ controls were compared against SARS-CoV-2 pseudoparticles or SARS-CoV-2 Spike expressing effector cells (see **Sup.Fig.3**). To validate our pDISPLAY approach cells were transfected with expression constructs for full length human ACE2 (hACE2 [FL]) or a human ACE2 where the signal peptide was replaced with the murine Ig κ-chain leader sequence (hACE2). In both instances the corresponding vector controls, pcDNA3.1 and pDISPLAY, were seperately transfected for comparison. The specificity of the SARS-CoV-2 and SARS-CoV assays were further confirmed by comparing hACE2-mediated fusion to human aminopeptidase N (hAPN) or dipeptidyl peptidase 4 (hDPP4) fusion, the coronavirus group I and MERS-CoV receptors, respectively. Lastly, in all assays target cells representing un-transfected cells (Mock) were also included. For pseudotype and cellcell fusion assays, luciferase assays were performed in duplicate and triplicate, respectively with the error bars denoting standard deviation.

**Supplemental Figure 3:**
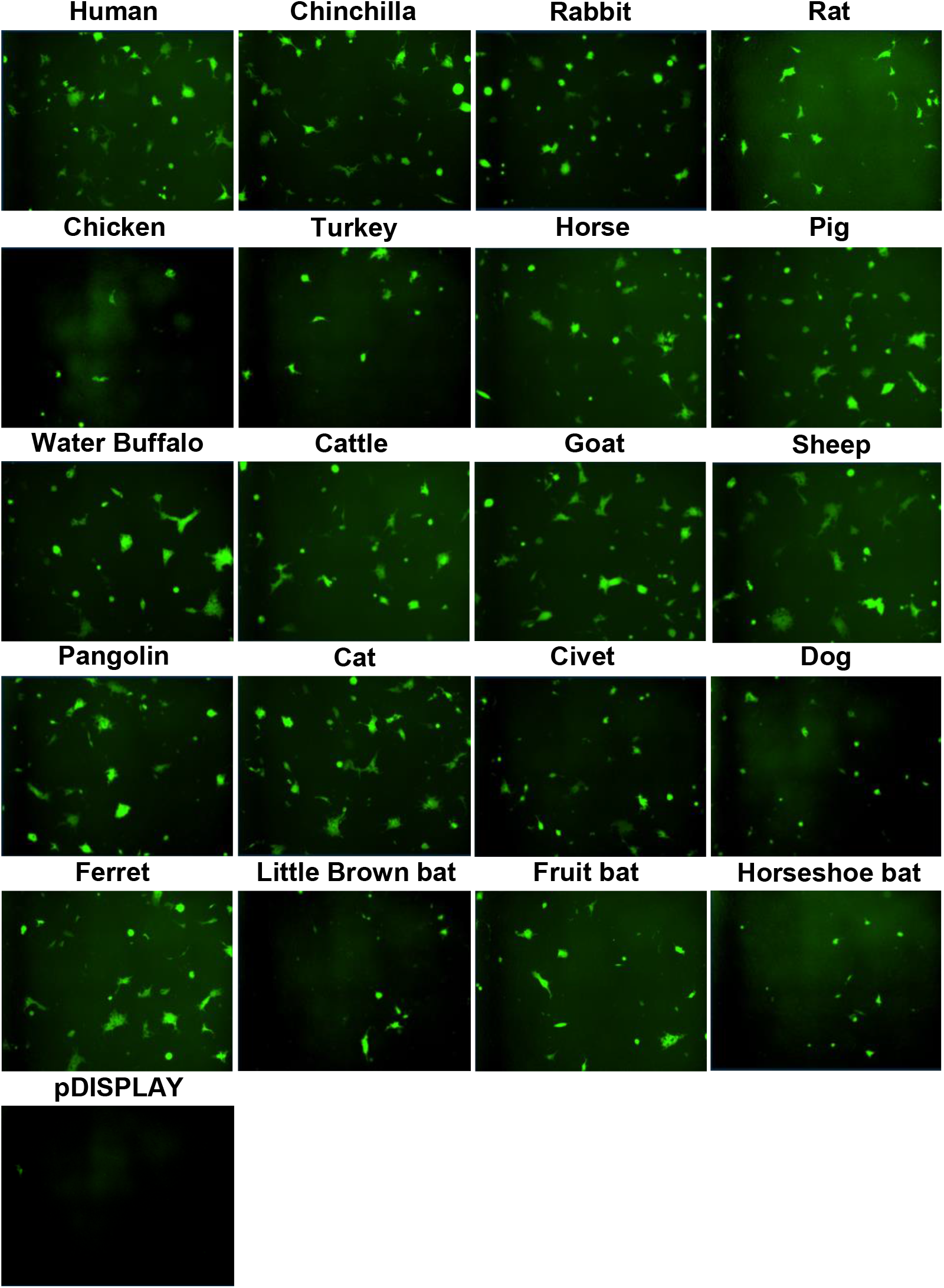
Syncytia formation following SARS-CoV-2 Spike expression. Effector cells expressing half of a split luciferase-GFP reporter and SARS-CoV-2 Spike were mixed with target cells expressing ACE2 proteins from the indicated hosts and the corresponding half of the reporter (see Methods). A vector only control was also included (pDISPLAY). Representative micrographs of GFP-positive syncytia formed following co-culturing are shown. Images were captured using an Incucyte live cell imager (Sartorius).

**Supplemental Figure 4:**
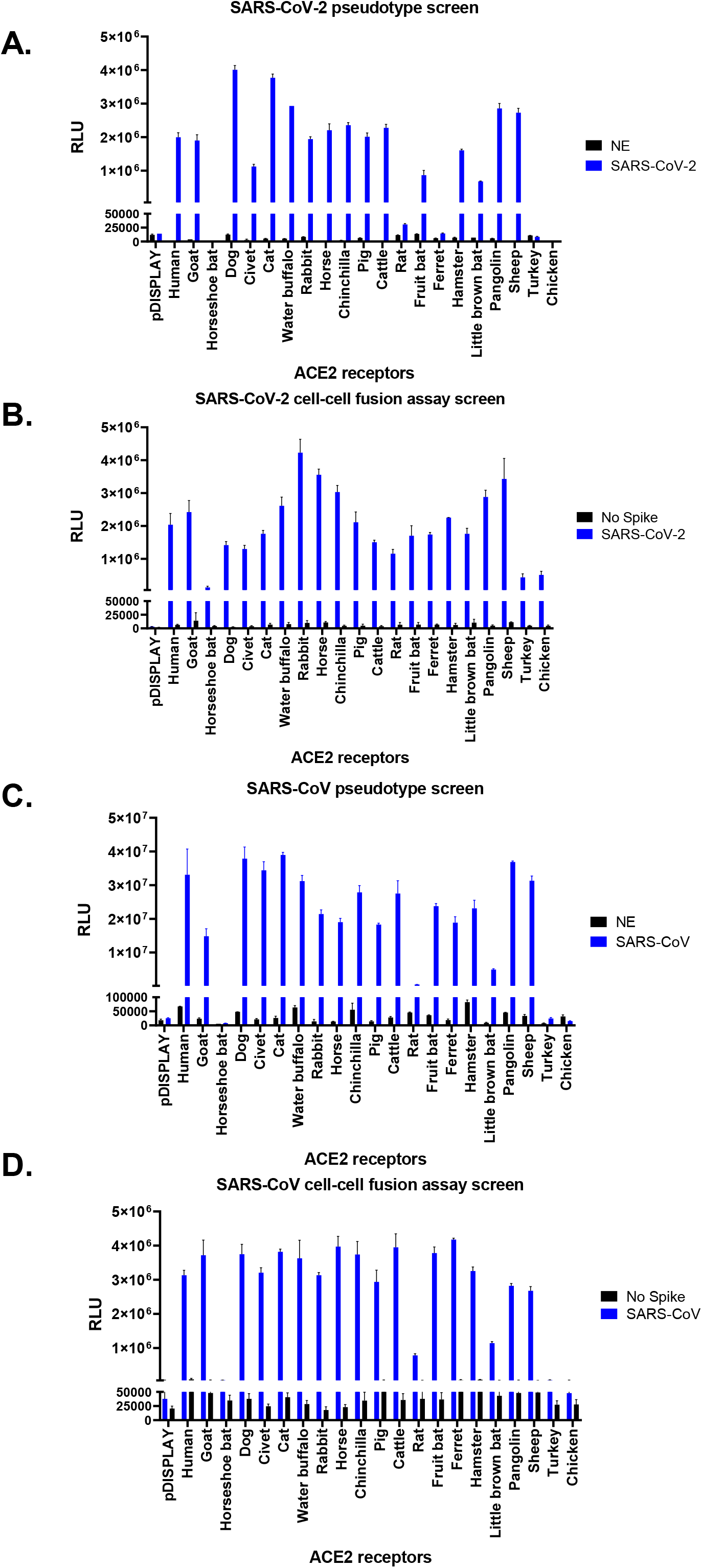
SARS-CoV-2 and SARS-CoV receptor usage screening. As per **Sup.Fig.2** NE and No Spike controls were included in all assays, as well as a vector only control (pDISPLAY). For pseudotype and cell-cell fusion assays, luciferase assays were performed in duplicate and triplicate, respectively with the error bars denoting standard deviation. Representative data sets from individual experiments are shown; however, the heat-maps and XY correlative plots in **Fig.2** and **Sup.Fig.5** summarise the results from three independent experiments performed on separate days.

**Supplemental Figure 5:**
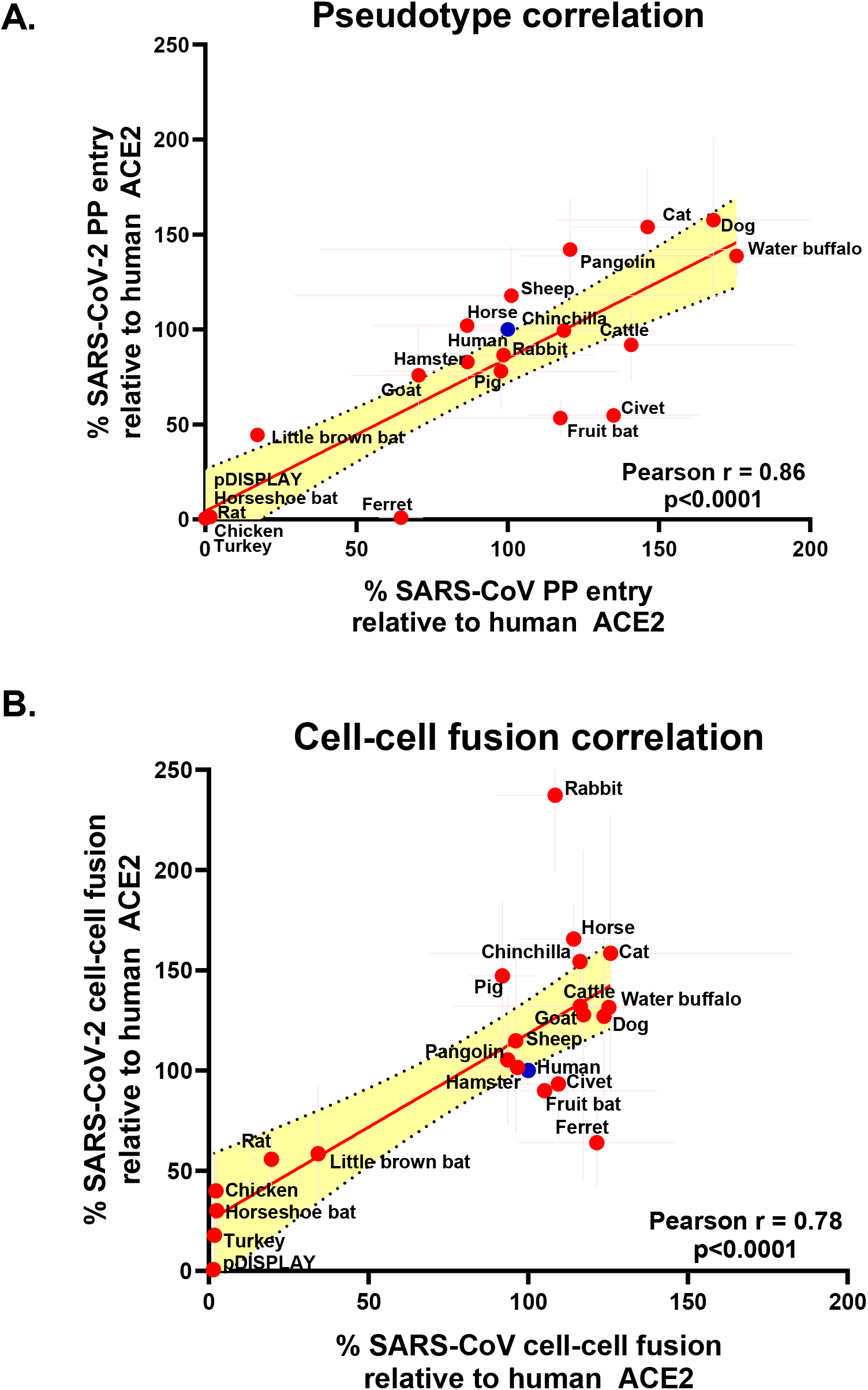
Correlating SARS-CoV-2 and SARS-CoV pseudotype and cell-cell fusion receptor usage. The receptor usage data for SARS-CoV-2 and SARS-CoV was examined by seperately comparing the pseudotype (**A**) or cell-cell fusion (**B**) assay results on XY scatter plots. The Pearson correlation was calculated and a linear line of regression fitted together with 95% confidence intervals. The x and y error bars denote the standard deviation from three experimental repeats performed on separate days. All values are plotted relative to the entry or cell-cell fusion recorded for human ACE2 (blue circles).

**Supplemental Figure 6:**
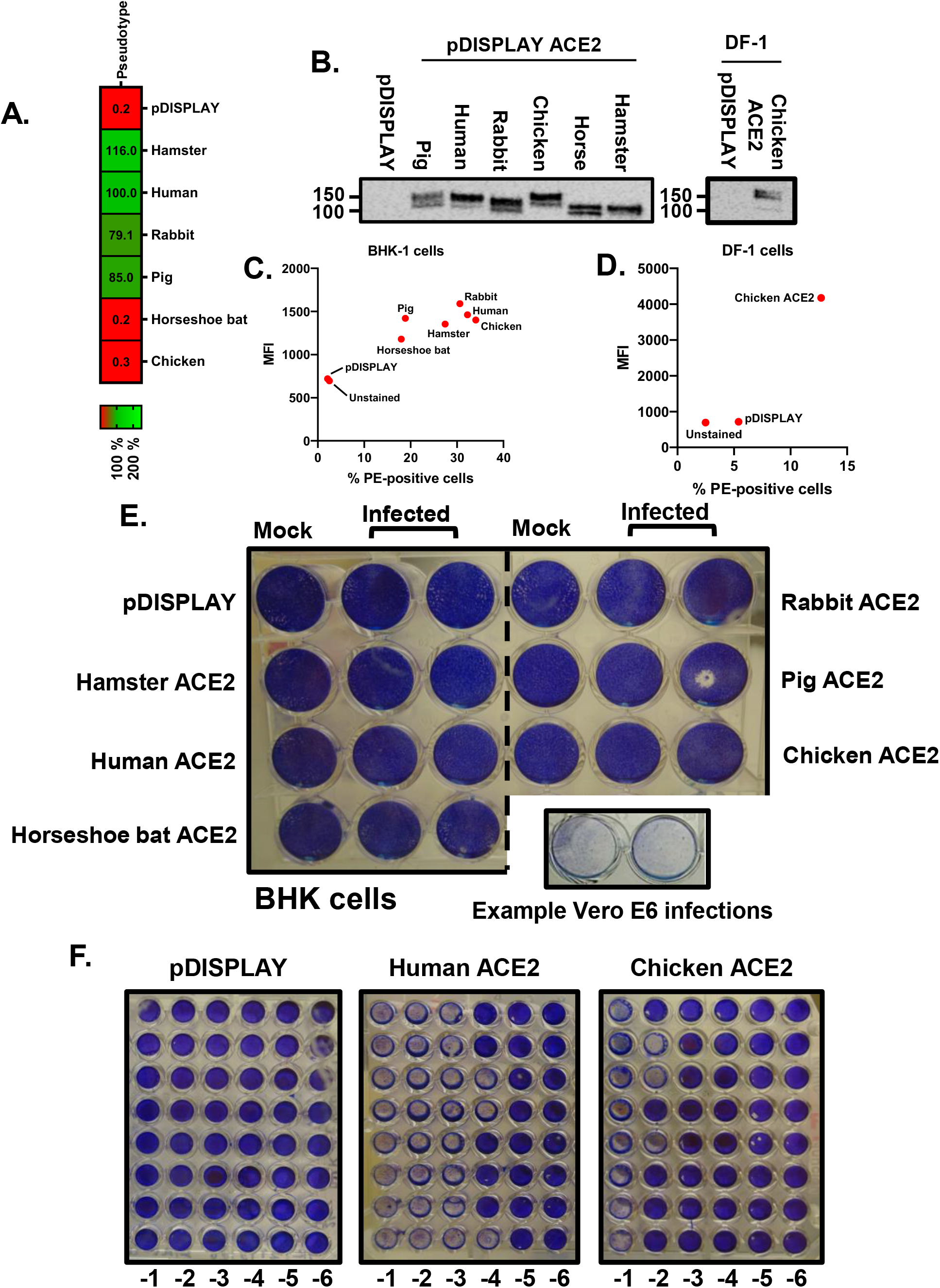
Experimental infection of cell lines over-expressing vertebrate ACE2 proteins. **(A)** SARS-CoV-2 pseudotype entry was assayed in BHK-21 transfected cells overexpressing ACE2 from the indicated species. Pseudotype infections were performed in triplicate and the mean value plotted on a heat map following normalisation to human ACE2. Similarly transfected target cells were lysed and the ACE2 expression analysed by western blot **(B)** or flow cytometry **(C)**. Equivalent experiments were performed for DF-1 cells **(B;** right panel, **D)**. **(E)** In parallel, BHK-21 cells were transfected with various ACE2-expression constructs and infected with SARS-CoV-2 at an MOI of 1. Cells were fixed and stained at 48 hpi. **(F)** Prior to fixation the supernatants from infected BHK-21 cells were removed for quantification of released virus by TCID-50. Representative images of these titrations, performed on Vero E6 cells, are shown (vector only control [pDISPLAY] as well as human and chicken ACE2).

**Supplemental Table 1:** Cell lines utilised in this study to quantify ACE2 mRNA levels and to assess virus permissibility.

| Cell line | Species | Organism | Cell type | Media and supplements |
| --- | --- | --- | --- | --- |
| MDCK | Canine | *Canis familiaris* | Kidney, epithelial | EMEM |
| D17 | Canine | *Canis familiaris* | Lung, epithelial | EMEM |
| NBL-2 | Canine | *Canis familiaris* | Kidney, epithelial | EMEM |
| LLC-RK1 | Rabbit | *Oryctolagus cuniculus* | Kidney, epithelial | Medium 199: horse serum, 1.12 g/L sodium bicarbonate |
| RK-13 | Rabbit | *Oryctolagus cuniculus* | Kidney, epithelial | EMEM |
| SIRC | Rabbit | *Oryctolagus cuniculus* | Cornea, fibroblast | EMEM |
| BHK-21 | Hamster | *Mesocricetus auratus* | Kidney, fibroblast | EMEM |
| CHO | Hamster | *Cricetulus griseus* | Ovary, epithelial-like | Hams F-12K: 20 mM HEPES |
| DEDE | Hamster | *Cricetulus griseus* | Lung, fibroblast | McCoy's 5a medium: 1.12 g/L sodium bicarbonate |
| NBL-6 | Horse | *Equus caballus* | Skin, fibroblast | EMEM |
| DF-1 | Chicken | *Gallus gallus* | Embryo, fibroblast | DMEM |
| LMH | Chicken | *Gallus gallus* | Liver, epithelial | Waymouth's medium |
| BT | Bovine | *Bos taurus* | Turbinate | DMEM |
| MDOK | Sheep | *Ovis aries* | Kidney, epithelial | EMEM |
| LLC-PK1 | Pig | *Sus scrofa* | Kidney, epithelial | Medium 199 |
| IPEC-J2 | Pig | *Sus scrofa* | Intestinal porcine enterocytes, epithelial | Hams F-12: 20 mM HEPES, 1% insulin/transferrin/selenium (ITS) |
| PK15 | Pig | *Sus scrofa* | Kidney, epithelial | EMEM |
| ST | Pig | *Sus scrofa* | Testis, fibroblast | EMEM |
| COS7 | Monkey | *Cercopithecus aethiops* | Kidney, fibroblast | DMEM |
| Vero E6 | Monkey | *Cercopithecus aethiops* | Kidney, epithelial | DMEM |
| Marc 145 | Monkey | *Cercopithecus aethiops* | Kidney, epithelial | DMEM |
| McCoy | Mouse | *Mus musculus* | Fibroblast | EMEM |
| NIH3T3 | Mouse | *Mus musculus* | Embryo, fibroblast | DMEM |
| Duck embryo fibroblast | Duck | *Anas platyrhynchos domesticus* | Embryo, fibroblast | EMEM |
| QT35 | Quail | *Coturnix coturnix* | Muscle, fibroblast | EMEM |

**Supplemental Table 2:**
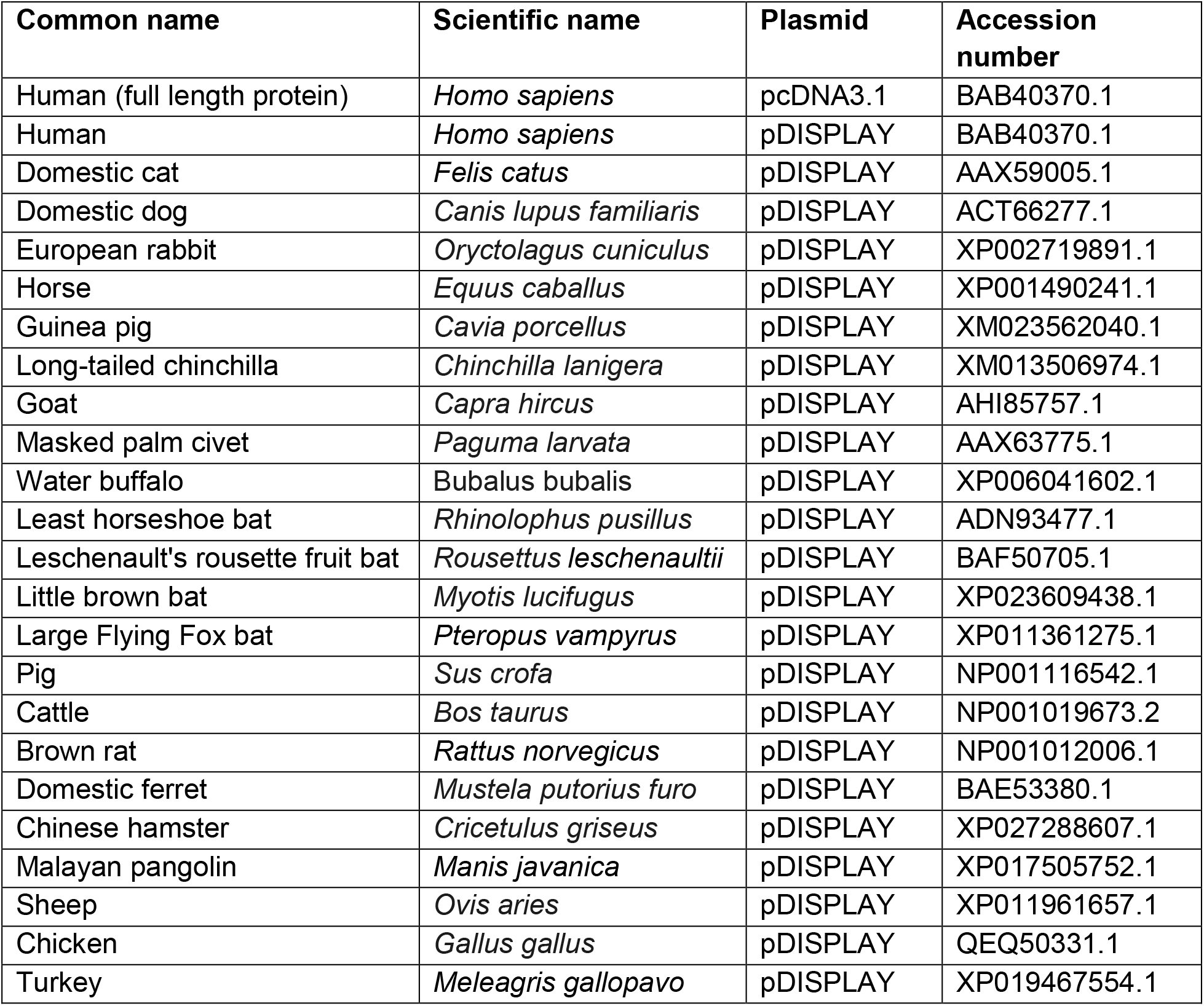
Codon optimised ACE2-expression plasmids used in this study for receptor usage screens.

**Supplemental Table 3:** β-coronavirus glycoproteins used in this study for receptor usage screens.

| Glycoprotein | Virus isolate | Backbone | Accession number | Source/Reference |
| --- | --- | --- | --- | --- |
| SARS-CoV-2 Spike | Wuhan-Hu-1 | pCAGGS | MN908947.3 | NIBSC, CFAR cat no. 100976 |
| SARS-CoV-1 Spike | ShanghaiQXC2 | pcDNA3.1+ with C-terminus FLAG tag | AAR86775.1 | Synthesised from NCBI sequence |

**Supplementary Table 4:**
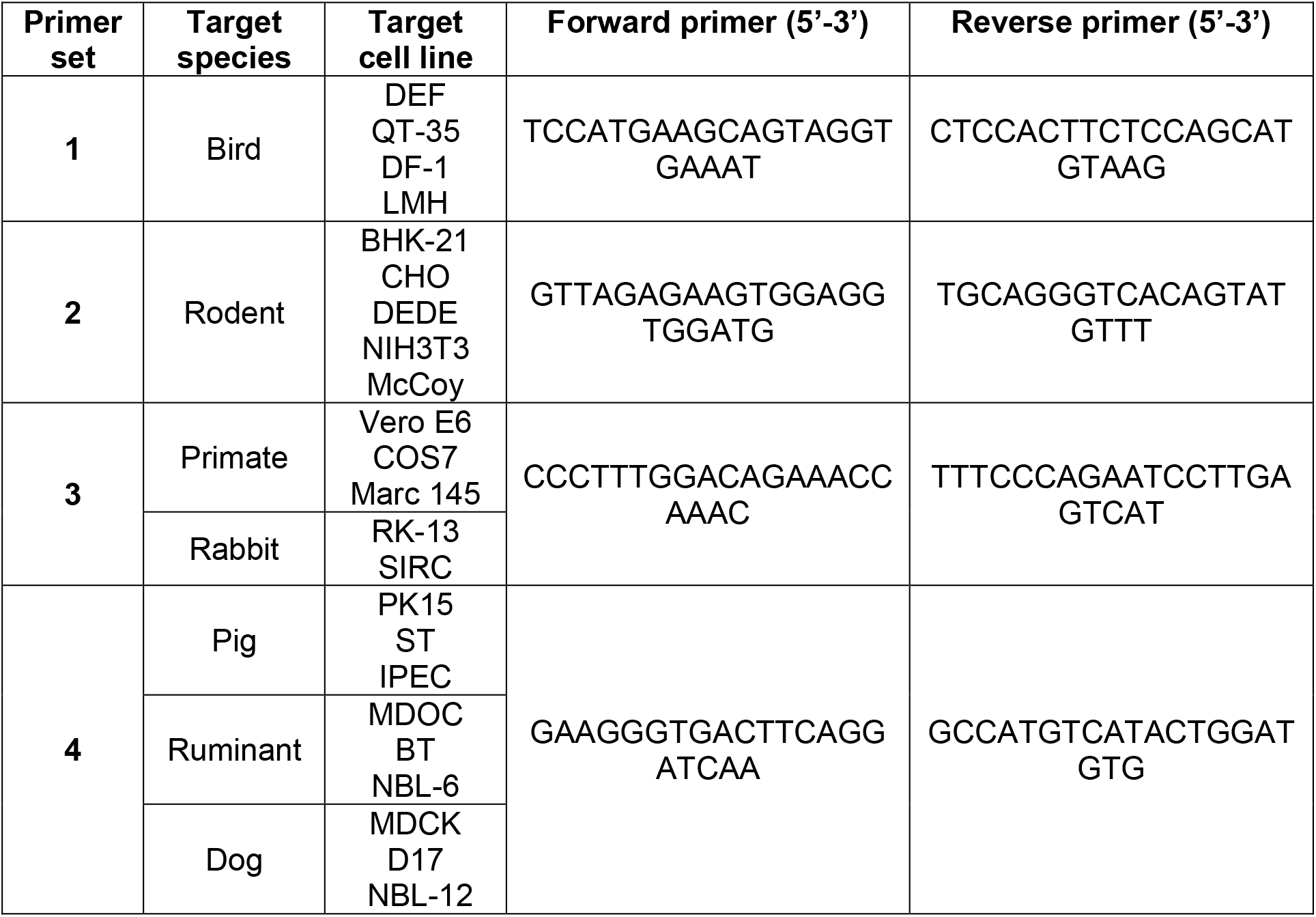
qPCR primer sets used in this study to quantify ACE2 mRNA levels.

## Notes

### Competing Interest Statement

The authors have declared no competing interest.

